# Differential role of dopamine D3 receptor through balanced modulation of Akt/mTOR and ERK_1/2_ activity in the reactivation of cocaine rewarding memories evoked by psychological versus physiological stress

**DOI:** 10.1101/2022.09.12.507553

**Authors:** Aurelio Franco-García, Rocío Guerrero-Bautista, Juana María Hidalgo, María Victoria Milanés, Victoria Gómez-Murcia, Cristina Núñez

## Abstract

Stress is an important trigger of relapses in cocaine use. These relapses engage the activity of memory-related nuclei, such as the basolateral amygdala (BLA) and the dentate gyrus (DG). Further, preclinical research signals D3 receptor (D3R) antagonists as promising therapeutic tools to attenuate cocaine reward and relapse. Therefore, we assessed the effect of SB-277011-A, a D3R antagonist, in the activity of Akt/mTOR and MEK/ERK_1/2_ pathways in these areas during the reinstatement of cocaine-induced conditioned place preference (CPP) evoked by psychological (restraint) and physiological (tail pinch) stress. Both stimuli reactivated the extinguished cocaine-CPP, but only restrained animals decreased their locomotor activity during reinstatement. Moreover, p-Akt, p-mTOR and p-ERK_1/2_ activity in the BLA and DG of restrained animals decreased during the reactivation of cocaine memories, contrasting to tail-pinched mice. While D3R blockade prevented stress-induced CPP reactivation and plasmatic corticosterone enhancement, SB-277011-A distinctly modulated Akt, mTOR and ERK_1/2_ activities in the BLA and DG based on the stressor and the dose of antagonist. Corticosterone may be partially responsible for these variations as we found high correlations among its levels and mTOR and/or Akt activity in the BLA and DG of restrained animals receiving SB-277011-A. Besides, locomotor activity of animals receiving 48 mg/kg of the antagonist highly correlated with p-mTOR/mTOR and p-ERK_1/2_ /ERK_1/2_ in the BLA during restraint- and tail pinch-induced relapse in cocaine-CPP, respectively. Hence, our study endorses D3R antagonists as therapeutic tools to prevent stress-induced relapses in drug use through a complex balance of Akt/mTOR and MEK/ERK_1/2_ pathways in memory-processing brain nuclei.

## 1. Introduction

Cocaine addiction is a cyclic condition in which addicts are unable to maintain prolonged periods of abstinence, thus eventuating in frequent relapses. The cocaine market has experienced a high growth in the last 5 years by improving the availability and quality of this drug, and therefore, attracting new customers. Hence, its consumption is a progressively major social problem due to the high prevalence of old and new users worldwide [1]

Environmental stimuli play a significant role in the transition to addiction to cocaine, in which an associative learning occurs [2]. Thus, memory formation and retrieval are essential in the development of this condition. Currently, therapeutic strategies are focused on relapse prevention, although clinical trials on multiple potential drugs to address this issue have been unsuccessful [3]. Extinction therapy tries to deal with this matter by creating new memories that inhibit abused drug-paired associations, but do not erase them. Nonetheless, this therapeutic approach has shown limited results, since previous drug-related memories can be reactivated by several triggers, such as environmental cues, the administration of a priming dose of the substance and stress [4, 5].

Whereas environmental cues and drug priming seem to provoke relapse by evoking cocaine-paired memories [6–8], different stressful episodes might not follow the same neural and biochemical networks that lead to similar behavioural outcome. The restraint and tail-pinch paradigms are robust models to induce physiological and psychological stress, respectively [4, 9]. Restraint models are based on the enclosure of rodents to make them experience a transient state of general anxiety [10]. On the other hand, tail-pinch consists of the application of mild non-traumatic pressure to the base of the rodent’s tail, which causes a light pain. Both paradigms have been extensively used to enhance the activation of the hypothalamic-pituitary-adrenal (HPA) axis and study drug-seeking related behaviours, as it has been previously reviewed [5].

Chronic exposure to some drugs of abuse is known to induce long alterations in the circuits critical for normal learning and memory processes. In the brain, ventral tegmental area (VTA) sends dopaminergic projections not only to the nucleus accumbens (NAcc) and prefrontal cortex, but interestingly to the basolateral amygdala (BLA), that is linked to emotional memory processes, and also to the hippocampus, in which the dentate gyrus (DG) plays a crucial role in episodic and spatial associations about rewarding and aversive events in the environment [11–13]. Moreover, ventral hippocampus acts as a place of integration between the spatial or contextual memory from the DG (dorsal hippocampus) and emotional information from the BLA [14]. Therefore, alterations in dopaminergic activity in the DG and BLA seem to work as modulators of drug-related memories. Concordantly, dopamine 3 receptor (D3R) has been proposed to play a large role in addiction-related behaviour, and therefore, to act as a potential target for relapse prevention in clinical therapy [15, 16]. Studies have shown the importance of D3R in relapse caused by stress, drug priming and drug cues [17, 18] but other findings provide conflicting results on the role of this receptor when its expression is suppressed [19, 20]. In addition, D3R blockade diminishes the availability of these receptors and, consequently, D3R-coupled biochemical pathways might be altered, as it has been published before [7, 8, 21]. D3R regulates the mitogen-activated protein kinase (MEK)-signal-regulated kinases (ERK), phosphatidyl inositol 3-kinase (PI3K)-protein kinase B (Akt) and mechanistic target of rapamicine (mTOR) intracellular signalling [21]. We have previously reported a beneficial effect of the antagonism of D3R to avoid drug-seeking reinstatement after psychosocial stress [6] but the impact of this antagonist on relapse in cocaine-seeking caused by other stressful stimuli remains yet to be elucidated. mTOR pathway has shown a major role in memory formation and recall through protein synthesis, as it has been reviewed before [22]. Since downregulation of this cascade has been linked to hippocampus neurite outgrowth [23] and enhancement of cognitive tasks [24], the precise control of this pathway remains conflicting. Our laboratory has previously reported an upregulation of mTOR pathway on BLA and DG after D3R blockade and psychosocial cocaine relapse prevention [8]. Similarly, cumulative evidence have reported the importance of MEK/ERK_1/2_ pathway in long term potentiation (LTP) and protein synthesis, critical events for memory formation and retrieval [25]. Hence, the aim of this work was to assess the impact of D3R blockade on the reinstatement of cocaine-seeking behaviour induced by psychological and physiological stress after the extinction of a previously acquired conditioned place preference (CPP) and elucidate the activity of Akt/mTOR and MEK/ERK_1/2_ pathways in the BLA and DG, and their possible relationship with the activity of the HPA axis during these events.

## 2. Materials and Methods

### 2.1 Animals

All animals (male C57BL/6 mice) were provided by Charles River laboratories (Saint-Germain-sur-l’Arbresle, France) at 6 weeks of age when they arrived at the laboratory and were stabled in groups of 4 in polypropylene cages (25L X 25W X 14.5H cm) in a room with controlled temperature (22 ± 2 ºC) and humidity (50 ± 10%), with ad libitum feeding and drinking (n= 73). Animals were adapted to a reversed 12 h light-dark cycle (lights off: 08:00 h – 20:00 h) for 7 days before the beginning of the experiments. Mice were handled for a week before the beginning of the experiment. All surgical and experimental procedures were performed in accordance with the European Communities Council Directive of 22 September 2010 (2919/63/UE) and were approved by the local Committes for animal research (Comité de Ética y Experimentación Animal; CEEA; RD 53/2013). Protocols were designed according to 3R to reduce the maximal number of mice and to minimise their suffering.

### 2.2 Drugs

Cocaine HCl (Alcaliber, Spain) was dissolved in sterile 0.9% NaCl. The antagonist D3R SB-277011-A (N-[trans-4-[2-(6-Cyano-3,4-dihydro-2(1H)-isoquinolinyl)ethyl]cyclohexyl]-4-quinolinecarboxamide dihydrochloride; Tocris, St. Louis, MO, USA) was dissolved in deionized distilled water (vehicle). All drugs were administered intraperitoneally (i.p.) in a volume of 0.01 ml/g body weight.

### 2.3 Conditioned Place Preference Paradigm

CPP paradigm was used to measure the rewarding properties of cocaine, as described previously [6]. The equipment used (Panlab, Barcelona, Spain) consisted of a box with two equally sized chambers (20 L × 18 W × 25 H cm) connected by a corridor (20L x 7W x 25H cm). The two larger chambers differed in their wall paint and floor texture and provided distinct contexts (visual and tactile cues). The walls of the corridor are transparent to reduce the time spent in this compartment. Manual guillotine doors were inserted during the conditioning sessions and removed during the tests. Weight transducer technology and PPCWIN software was used to detect and analyse animal position and number of entries in each compartment during the test.

The CPP paradigm consists of three different phases: a preconditioning phase, a conditioning phase, and a testing phase. Firstly, during the pre-conditioning phase (Pre-C), animals freely explored all compartments for 15 min and the time spent in each compartment was analysed. Those showing a natural preference (> 67%) or aversion (< 33%) for any of the compartments were excluded from the experiment. All the behavioural experiments were carried out at the same time of the day.

Mice were counterbalanced for compartment assignment in terms of initial spontaneous preference. A Student’s *t*-test was used to confirm that there were no significant differences in term of time spent in compartment between groups (in cocaine-paired and the saline-paired compartments) during Pre-C. The following day, during the conditioning phase, doors were closed to avoid animals pass to other chambers.

All animals were conditioned with 25 mg/kg of cocaine on days 1 and 3 during 30 min. After that, they went back to the home cage and 4 h later they received a saline injection before being conditioned to the vehicle-paired compartment for another 30 min. 25 mg/kg of cocaine dose has been previously shown to be adequate to induce a strong CPP [6–8, 26]. On days 2 and 4, mice received firstly a saline injection before being confined to the vehicle-paired chamber and 4 h after placing them to the home cage, animals were injected with cocaine before confinement in the drug-paired chamber for another 30 min. After that, on day 5, a post-conditioning (Post-C) test was carried out exactly as Pre-C test, animals exploring all compartments during 15 min. On day 6 and for the following 7-8 weeks animals underwent extinction sessions twice a week in no consecutive days. During these sessions mice explored freely all the compartments during 15 min as in Pre-C and Post-C tests. The criterion for the extinction of drug seeking behaviour was considered when there were no significant differences (Student’s *t*-test) in the time spent by each group in the cocaine-associated chamber during the ext test with regard to that in the Pre-C test. To validate the extinction, a new session was repeated 48h later.

To study the effect of psychological or physiological acute stress on the reinstatement of cocaine-induce CPP, 2 days after extinction was confirmed, animals were subjected to restraint or tail pinch, respectively, followed by a reinstatement (reinst) test to analyze the relapse in cocaine-induced CPP. In this reinst test, mice were placed in the central corridor of the CPP apparatus during 15 min and allowed to explore freely all the compartments (as in pre-C, post-C, and ext tests).

All the stressful episodes were carried out in a different room where the CPP apparatus was placed. To induce restraint stress, mice were placed for 15 min in cylindrical plastic restrainers (2.5 cm diameter X 11.5 cm length) with an aperture in one of the ends to allow normal air flow. After that, mice passed the reinst test. For the tail pinch stress, another group of animals were put in a plastic cage (25 H X 25 W X 14.5 cm) and a binder clip (7 mm wide, with oval inner section of 8 mm wide, and 13 mm height) was place in the last third of the tip of the tail for 15 minutes (clamping force, 10 newton). After that, mice were place in the CPP apparatus to pass the reinst test. For each stressor, a group of control animals were put in a plastic cage (25 H X 25 W X 14.5 cm) for 15 min without being subjected to any stressful stimulus for 15 minutes and then were tested for reinstatement of cocaine-CPP.

The role of D3R in the reactivation of cocaine-induced CPP was investigated by the administration of a single dose of the D3R antagonist, SB-277011-A (24 or 48 mg/kg) or its vehicle to mice 30 min before the stressful session.

Test data (time spent in each chamber of the apparatus and number of crossings between them) were recorded automatically by PPCWIN software (Panlab, Barcelona, Spain). As these data were collected by a computer, blinding to experimental group was not required.

### 2.4 Corticosterone measurements

After behavioural experiments, mice were sacrificed by cervical dislocation, trunk blood samples were collected and to separate the plasma they were centrifuged at 960 g 15 min, 4 °C. Samples were stored at −80 °C until their analysis. Corticosterone levels were determined using commercially available kits for mice (125 I-corticosterone radioimmunoassay; MP Biomedicals, USA). The sensitivity of the assay was 7.7 ng/ml.

### 2.5 Tissue Collection and Western Blot Analysis

Brains were removed and placed at −80 °C until they were processed. Brain interest regions were cut in 500 µm coronal slides on a cryostat at −20 °C and stored at −80 °C. BLA and DG were cut in two consecutive slides corresponding approximately −1.47 to 2.19 mm from Bregma for DG and −0.95 to −1.59 mm for BLA according to the atlas of Franklin and Paxinos (2007) [27]. Bilateral 1 mm^2^ punches of the DG and BLA were stored into tubes containing 50 µL of homogenization solution (with phosphate buffered saline (PBS), sodium lauryl sulfate, proteases inhibitors, and a phosphatases inhibitors cocktail set), as previously described by Leng et al (2004) [28]. Tubes containing punches were consequently frozen in dry ice and place at −80 °C until processing. Samples were sonicated, vortexed, and sonicated again before centrifugation at 11000 g for 10 min at 4 °C; then, the supernatant was recovered and placed at −80°C. BCA method was used for protein determination. Twenty µg of protein samples were loaded and separated on 4–12% Criterion Bis-Tris Gels (Millipore, USA) electrophoresis and transferred to polyvinylidene fluoride (PVDF) membranes (EMD Millipore Corporation, USA). Membranes were saturated with 1% BSA in TBST (tris buffer saline tween, 0.15%) and respectively incubated overnight with primary and corresponding secondary antibodies [rabbit anti p-mTOR (1:1000; #5536, Cell Signaling Technology, Danvers, MA, USA); rabbit anti mTOR (1:1000; #2983, Cell Signaling Technology), rabbit anti p-Akt (1:750; #4060L, Cell Signaling Technology), rabbit anti Akt (1:1000; #9272s, Cell Signaling Technology), mouse anti p-ERK_1/2_ (1:750; #sc-7383, Santa Cruz), rabbit anti ERK2 (1:1000; sc-154, Santa Cruz), anti-rabbit antibody (1:10000; #31430; Thermo Fisher Scientific, Waltham, MA, USA), anti-mouse antibody (1:10000; #31460; Invitrogen)]. Immunoreactivity was detected with an enhance chemiluminescent Western Blot detection system (ECL Plus, Thermo Fisher Scientific) and visualised by an Amersham Imager 680 (General Electric, Boston, MA, USA). Antibodies were stripped from the blots by incubation with stripping buffer (glycine, 25 mM and SDS 1%, pH 2) for 30 min at 37 °C. Results were normalised to β-actin, and quantifications were performed using ImageQuant TL 8.1 software (General Electric). The ratio of p-Akt/β-actin, Akt/β-actin, p-mTOR/β-actin, mTOR/β-actin, p-ERK_1/2_/β-actin and p-Akt/Akt, p-mTOR/mTOR, p-ERK_1/2_/ERK were plotted and analysed. Protein levels were corrected for individual levels.

### 2.6 Statistical Analysis

All descriptive data were presented as means and standard error of means (S.E.M). To consider the extinction in behavioural experiments, a paired Student’s *t*-test was performed to analyse the difference between the time spent by mice in the cocaine-paired compartment during the pre-C and post-C tests. For the time spent in the cocaine-paired chamber, the statistical analysis was performed using one-way ANOVA with repeated measures followed by the Tukey multiple comparisons test to determine specific group differences. Number of entries during ext and reinst tests were analysed through a paired Student’s *t* test. The reinst score, plasma corticosterone levels and western blot data were analysed using one-way ANOVA followed by the Tukey *post-hoc* test. Correlations between different parameters were assessed using the Pearson’s correlation coefficient. Differences with a P < 0.05 were considered significant. Statistical analyses were performed with GraphPad Prism 9 (GraphPad Software Inc., San Diego, CA, USA).

## 3. Results

### 3.1 D3R blockade impeded the reactivation of cocaine-induced CPP induced by both psychological and physiological stress

It is broadly known that cocaine conditioning provokes a strong CPP in rodents [29]. Accordingly, all the mice used in this work spent significantly more time in the cocaine-associated compartment during the post-C test regarding the pre-C test and, after the extinction sessions, the period that animals stayed in the drug-paired chamber in the ext test was statistically similar to that in the pre-C test (Fig. 1B,D). We used two types of stressors, one psychological – restraint – and one physiological – tail pinch –, to induce the reactivation of the cocaine-associated rewarding memories responsible for the triggering of drug-seeking behaviours. In contrast to control mice, that were not subjected to any stressful stimuli, the animals that received vehicle instead the D3R antagonist before an acute session of restraint or tail pinch spent significantly more time in the drug-paired chamber during the reinst test with regard to that in the ext and pre-C tests (Fig. 1B,D). Additionally, as we had previously reported [6], the antagonism of D3R with 48 mg/kg but not 24 mg/kg of SB-277011-A, prevented the significant enhancement of the time spent by mice in the cocaine-associated compartment during the reinstatement test evoked by any of these two stressors (Fig. 1B,D). When we calculated the reinstatement score (as the difference of time that mice spent in the cocaine-paired compartment during the reinst test minus that in the ext test), we found a significant augment in animals pre-treated with vehicle before the stressful episode regarding controls, and only the higher dose of the D3R antagonist was able to significantly block this effect (Fig. 1C,E).

**Figure 1.**
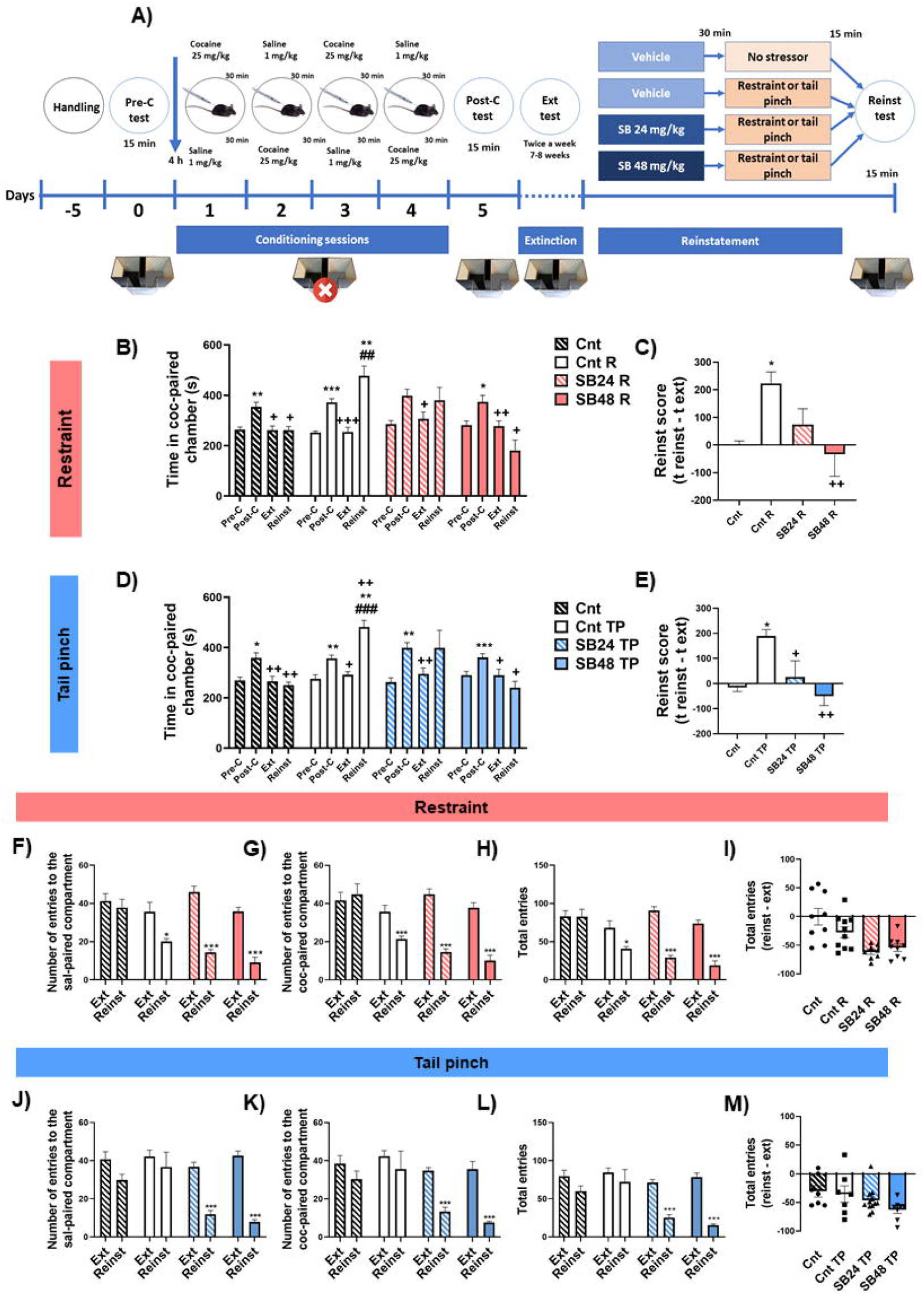
**The antagonism of D3R blocked the reactivation of cocaine-induced CPP induced by both psychological and physiological stress. (A) Schematic of the experimental timeline and behavioural procedures**. (**B) Mean preference time spent in cocaine-paired chamber during Pre-C, post-C, Ext and Reinst of control mice (vehicle, no stressor; Cnt), animals that experienced restraint but did not receive the D3R antagonist (vehicle, restraint; Cnt R), and animals that experienced restraint and received the D3R antagonist at two different doses, 24 or 48 mg/kg, restraint; SB24 R or SB48 R)**. Repeated measures one-way ANOVA showed differences among tests within Cnt group (F (2.182, 17.46) = 12.86; P = 0.0003), Cnt R group (F (1.461, 13.15) = 23.59; P = 0.0001), SB48 R (F (1.435, 12.91) = 7.776; P = 0.0098) but not SB24R group (F (1.432, 10.02) = 3.418; P = 0.0845). ^*^p < 0.05, ^**^p < 0.01, ^***^p < 0.001 vs Pre-C; ^+^p < 0.05, ^++^p < 0.01, ^+++^p < 0.001 vs Post-C; ^##^p < 0.01 vs Ext (Tukey’s test). **(C)Reinstatement score expressed as time in cocaine-paired chamber during Reinst test minus the same in Ext test in Cnt, Cnt R, SB24 R and SB48 R animals**. One-way ANOVA revealed significant differences among group means (F (3, 32) = 4.913; P = 0.0064). ^*^p < 0.05 vs Cnt; ^++^p < 0.01 vs Cnt R (Tukey’s test). (**D) Mean preference time spent in cocaine-paired chamber during Pre-C, post-C, Ext and Reinst of control mice (vehicle, no stressor; Cnt), animals that experienced tail pinch but did not receive the D3R antagonist (vehicle, tail pinch; Cnt TP), and animals that experienced tail pinch and received the D3R antagonist at two different doses (D3R antagonist at SB-277011-B, 24 or 48 mg/kg, restraint; SB24 TP or SB48 TP)**. Repeated measures one-way ANOVA showed differences among tests within Cnt group (F (2.440, 17.08) = 14.24; P = 0.0001), Cnt TP group (F (1.917, 15.34) = 30.18; P < 0.0001), SB48 TP group (F (1.644, 11.51) = 6.606; P = 0.0154) but not SB24 TP group (F (1.314, 14.45) = 3.561; P = 0.0706). ^*^p < 0.05, ^**^p < 0.01, ^***^p < 0.001 vs Pre-C; ^+^p < 0.05, ^++^p < 0.01 vs Post-C; ^###^p < 0.001 vs Ext (Tukey’s test). **(E) Reinstatement score expressed as time in cocaine-paired chamber during Reinst test minus the same time in Ext test in Cnt, Cnt TP, SB24 TP and SB48 TP animals**. One-way ANOVA revealed significant differences among group means (F (3, 30) = 5.505; P = 0.0039). ^*^p < 0.05 vs Cnt; ^++^p < 0.01 vs Cnt TP (Tukey’s test). **(F-H) Locomotor activity during Ext and Reinst tests for Cnt, Cnt R, SB24 R and SB48 R animals**. Graphics show the number of entries to saline-paired (paired Student’s *t* test: Cnt (t_8_ = 0.5483, P = 0.5985), Cnt R (t_9_ = 2.930, P = 0.0167), SB24 R (t_7_ =10.74, P < 0.0001) and SB48 R (t8 = 8.321, P < 0.0001)) and cocaine-paired (paired Student’s *t* test: Cnt (t_8_ = 0.4024, P = 0.6979), Cnt R (t_8_ = 3.608, P = 0.0069), SB24 R (t_7_ = 11.87, P < 0.0001) and SB48 R (t_8_ = 8.168, P < 0.0001)) compartments and total entries (paired Student’s *t* test: Cnt (t_8_ = 0.02346, P = 0.9819), Cnt R (t_9_ = 2.878, P = 0.0182), SB24 R (t_7_ = 14.27, P < 0.0001) and SB48 R (t_8_ = 8.602, P < 0.0001)) **(H)**. ^*^p < 0.05, ^***^p < 0.001 vs Ext test (Tukey’s test). **(I) Total entries score in Cnt, Cnt R, SB24 R and SB48 R animals, expressed as the difference in total entries during Reinst test minus total entries during Ext test**. One-way ANOVA: (F (3, 32) = 8.240; P = 0.0003). **(J-L) Locomotor activity during Ext and Reinst tests for Cnt, Cnt TP, SB24 TP and SB48 TP animals**. Graphics show the number of entries to saline (sal)-paired (paired Student’s *t* test: Cnt (t_7_= 1.619, P = 0.1495), Cnt TP (t_8_ = 0.5979, P = 0.5664), SB24 TP (t_11_ = 7.794, P < 0.0001) and SB48 TP (t_7_ = 14.24, P < 0.001) **(J)**, cocaine (coc)-paired (paired Student’s *t* test: Cnt (t_7_ = 1.015, P = 0.3438), Cnt TP (t_8_ = 0.5979, P = 0.5377), SB24 TP (t_11_ = 6.390, P < 0.0001) and SB48 TP (t7 = 6.312, P = 0.0004) **(K)** compartments and total entries (paired Student’s *t* test: Cnt (t_7_ = 1.405, P = 0.2028), Cnt TP (t_8_ = 0.6641, P = 0.5253), SB24 TP (t_11_ = 7.386, P < 0.0001) and SB48 TP (t_7_ = 10.19, P < 0.0001) **(L)**. ^*^p < 0.05, ^***^p < 0.001 vs Ext test. **(M) Total entries score in Cnt, Cnt TP, SB24 TP and SB48 TP animals, expressed as the difference in total entries during Reinst test minus total entries during Ext test**. One-way ANOVA: (F (3, 30) = 2.297; P = 0.0977). Data are shown as mean ± S.E.M (n = 7 – 12 per group).

### 3.2. The locomotor activity of mice during the reactivation of cocaine-induced CPP is dependent on the kind of triggering stressor

Furthermore, we intended to study the locomotor activity of mice during the reactivation of cocaine-associated memories induced by acute psychological or physiological stress. To accomplish that, we analysed the number of entries to the cocaine- and saline-paired chambers and the sum of all of them during the reinst and ext tests. Whereas there were no changes in the number of total entries as well as those to any of the compartments in control animals during both reinst and ext tests, mice that were restrained significantly diminished their entries to any of the compartments of the CPP apparatus as well as the total entries during the reinst test when compared with the ext test (Fig. 1F-H). Nonetheless, this effect was not observed in mice that reinstated the cocaine-CPP after an episode of tail pinch (Fig. 1I-K). In addition, after the administration of both doses −24 and 48 mg/kg - of SB-277011-B we observed a significant decrease in the activity of mice subjected to any of the stressors during the reinst test regarding the ext test (Fig. 1F-H; J-L). To investigate whether D3R antagonism influenced the altered locomotor activity during the reactivation of cocaine-associated memories induced by restraint or tail pinch, we analysed the difference between the number of total entries in the reinst test and that in the ext test for each experimental group. However, there were no significant changes between animals pre-treated with vehicle before any of the stressful episodes and the mice injected with the D3R antagonist prior being subjected to restraint or tail pinch (Fig. 1I,M).

### 3.3 Corticosterone plasma levels might influence the reinstatement of cocaine-CPP after physiological but not psychological stress

To investigate whether the hormonal response to stress affects the reinstatement of cocaine-CPP, we quantified corticosterone plasma concentration after the reinst test in all the experimental groups. Similar to our previous work [6], we found that after the reactivation of cocaine-associated rewarding memories evoked by both restraint or tail pinch the levels of corticosterone in plasma significantly augmented regarding control animals, and that the administration of both 24 and 48 mg/kg of the D3R antagonist blocked this enhancement (Fig. 2A,B). Curiously, we detected that corticosterone concentration of restrained mice was significantly higher in comparison with animals that were subjected to tail pinch to induce the reactivation of cocaine-CPP (Fig. 2C). Moreover, corticosterone plasma concentration highly correlated positively with the reinstatement score of animals that received vehicle and were subjected to tail pinch to induce the reactivation of cocaine rewarding memories, while this correlation was not significant in mice that were injected with vehicle and restrained (Fig. 2D,E). In addition, there were not significant correlations between the reinstatement score of the animals injected with both doses of SB-277011-A prior the tail pinch or restraint episode and their corticosterone plasma levels (Fig. 2D,E), thus suggesting that D3R would modulate this association at least when the relapse in cocaine-CPP is triggered by a physiological stressor. Besides, corticosterone plasma levels moderately correlated positively with the difference of total entries between the reinst and ext tests of mice receiving D3R antagonist or its vehicle before being subjected to a psychological or physiological stressor to induce the relapse in cocaine-CPP (Fig. 2F,G). However this difference of entries did not correlate with the reinstatement score of mice restrained or tail pinched after being administered with vehicle or SB-277011-A (Fig 2H,I). Collectively, these data appear to signal that, although stress could augment the locomotor activity of animals, this activity is unlikely to be decisive for the reinstatement of cocaine-seeking behaviours.

**Figure 2.**
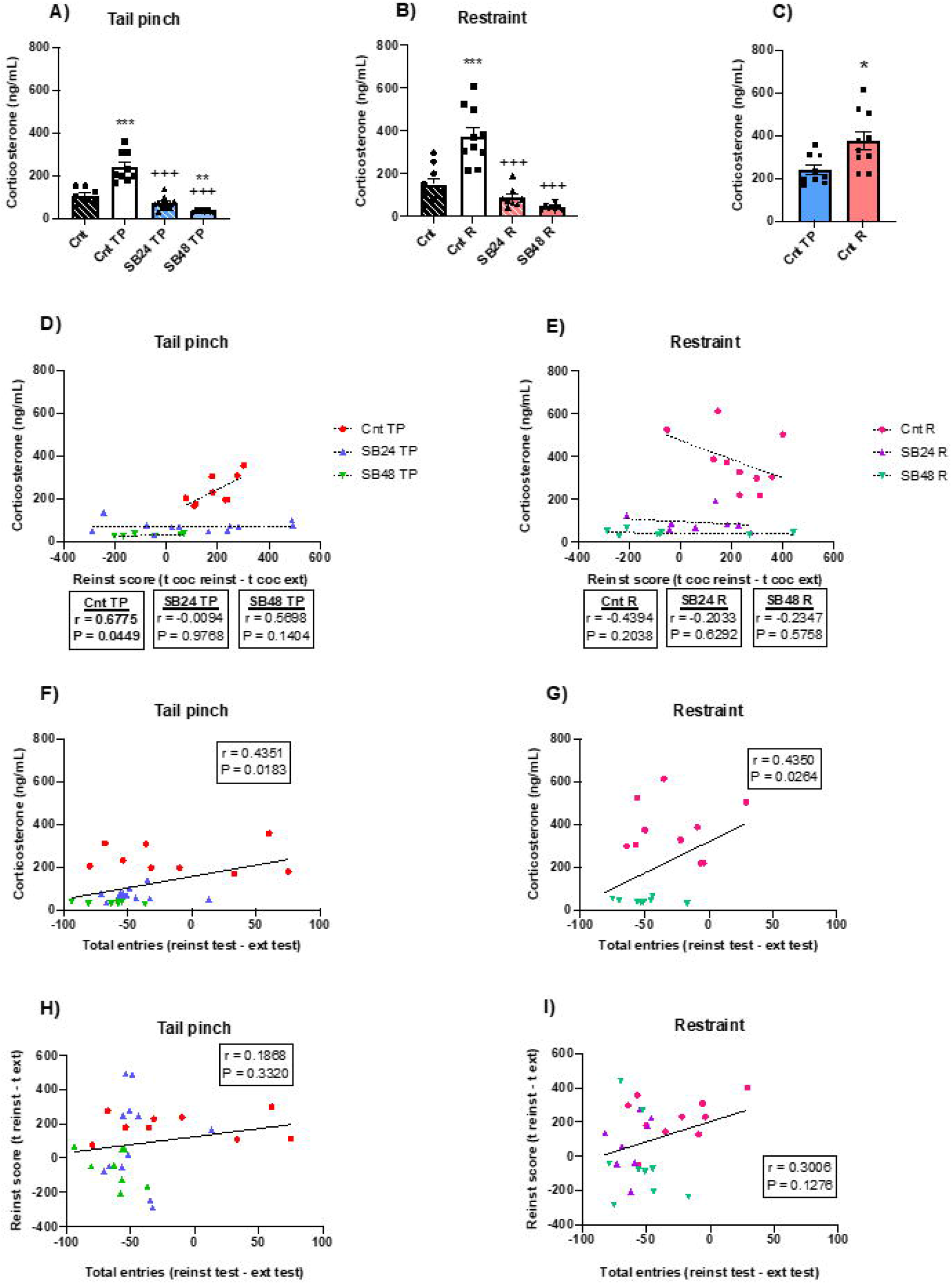
**(A) Analysis of corticosterone levels in plasma of control mice (vehicle, no stressor; Cnt), animals that experienced tail pinch but did not receive the D3R antagonist (vehicle, tail pinch; Cnt TP), and animals that experienced tail pinch and received the D3R antagonist at two different doses (SB-277011-B, 24 or 48 mg/kg, tail pinch; SB24 TP or SB48 TP)**. One-way ANOVA: F (3, 32) = 42.59); P < 0.001. ^**^p < 0.01, ^***^p < 0.001 vs Cnt, ^+++^p < 0.001 vs Cnt TP (Tukey’s test). **(B) Analysis of corticosterone levels in plasma of control mice (vehicle, no stressor; Cnt), animals that experienced tail pinch but did not receive the D3R antagonist (vehicle, restraint; Cnt R), and animals that experienced restraint and received the D3R antagonist at two different doses SB-277011-B**,**24 or 48 mg/kg, restraint; SB24 R or SB48 R)**. One-way ANOVA: F (3, 31) = 27.29; P < 0.001. ^***^p < 0.001 vs Cnt, ^+++^p < 0.001 vs Cnt R (Tukey’s test). **(C) Comparison of corticosterone levels in plasma between Cnt TP and Cnt R groups**. Unpaired Student’s *t* test: t_17_ = 2.809, P = 0.0121. ^*^p < 0.05 vs Cnt TP. **(D) Correlation analysis between plasmatic corticosterone and reinstatement (reinst) score in tail pinched animals, measured as the time spent in cocaine-paired chamber during reinst test minus the same during extinction (ext) test. (E) Correlation analysis between plasmatic corticosterone and reinstatement (reinst) score in restrained animals. (F) Correlation analysis between plasmatic corticosterone and total entries in tail pinched animals, measured as the total entries during reinst test minus those during ext test. (G) Correlation analysis between plasmatic corticosterone and total entries in restrained animals. (H) Correlation analysis between reinst score and total entries in tail pinched animals. (I) Correlation analysis between reinst score and total entries in restrained animals**. Correlation analyses were revealed through Pearson’s test. Data are shown as mean ± S.E.M (n = 7 – 12 per group).

### 3.4 The type of stressor affects differently to the activity of Akt/mTOR pathway and ERK_1/2_ in the BLA and DG of mice that relapsed in cocaine-CPP

We next studied the activity, as the ratio between the phosphorylated (p)- and total protein, of ERK_1/2_, Akt and mTOR since they have been long related to the plastic changes that occur during the formation and retrieval of different kind of memories [22, 25] and are regulated by D3R [7, 8, 21], in the BLA and the DG.

In the BLA of mice that were restrained in order to reactivate the cocaine-induced CPP we uncovered significant decreases regarding controls in the levels of p-Akt and p-mTOR and a clear trend to diminish of p-ERK_1/2_ (Fig. 3A,D,G) that paralleled with invariable amounts of total Akt, mTOR and ERK_1/2_ (Fig. 3B,E,H), thus leaded to diminished p-Akt/Akt, p-mTOR/mTOR and p-ERK_1/2_/ERK_1/2_ ratios, although the last was not statistically significant (Fig. 3C,F,I). On the contrary, in the BLA of mice subjected to tail pinch before the reinst test we did not observed alterations in the levels of p-Akt, p-mTOR or p-ERK_1/2_ (Fig. 3J,M,P) nor their total protein levels (Fig. 3 K,N,Q), hence the activity of these kinases did not differ between experimental groups (Fig. 3L,O,R). Similarly, in the DG of restrained animals prior the reinst test we found significant decreases in the levels of p-Akt, p-mTOR and p-ERK_1/2_ in comparison with control animals (Fig. 4A,D,G). Despite significant lower levels of total Akt, but not of mTOR nor ERK_1/2_ (Fig. 4B,E,H), the ratios p-Akt/Akt, p-mTOR/mTOR and p-ERK_1/2_/ERK_1/2_ in the DG after the reinstatement of cocaine-CPP induced by restraint were diminished with regard to control group, although only p-ERK_1/2_/ERK_1/2_ reached statistical significance (Fig. 4C,F,I). On the contrary, animals subjected to an acute session of tail pinch to induce the relapse in cocaine-CPP did not exhibit alterations in p-Akt, p-mTOR nor p-ERK_1/2_ levels in the DG regarding controls (Fig. 4J,M,P). Also, no differences in total Akt, mTOR and ERK_1/2_ between Cnt and Cnt-TP groups were detected (Fig. 4K,N,O). As a result, p-Akt/Akt, p-mTOR/mTOR and p-ERK_1/2_/ERK_1/2_ did not significantly change in this hippocampal region after the reinstatement of cocaine-CPP evoked by a physiological stressor when compared with controls (Fig. 4L,O,R).

**Figure 3.**
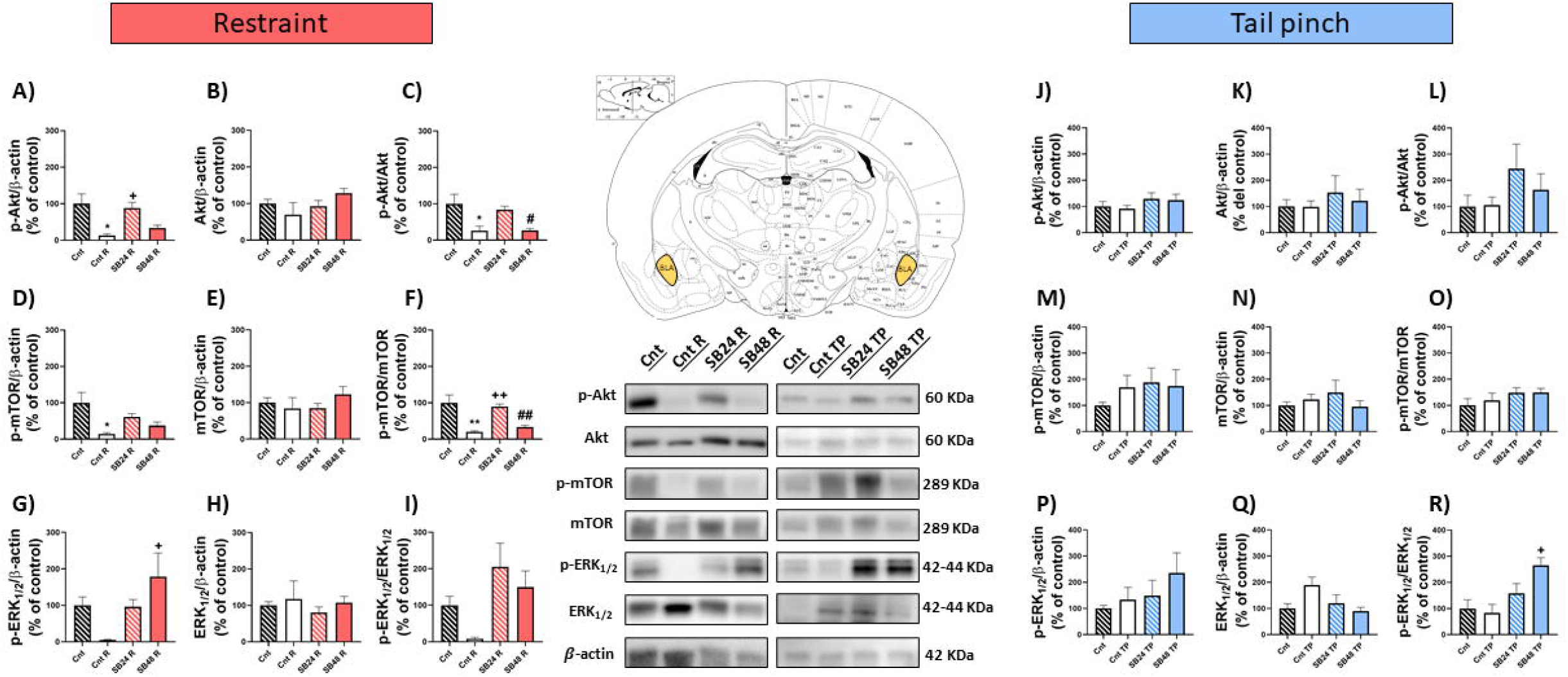
**Basolateral amygdala analysis in restraint and tail pinch stress paradigms. (A-C) Semiquantitative analysis and representative immunoblot of p-Akt/β-actin, Akt/β-actin and p-Akt/Akt levels in restraint paradigm**. One-way ANOVA: F (3, 21) = 5.454; P = 0.0062 **(A)**; F (3, 21) = 1.774; P = 0.1829 **(B)**; F (3, 21) = 6.916 **(C). (D-F) Semiquantitative analysis and representative immunoblot of p-mTOR/β-actin, mTOR/β-actin and p-mTOR/mTOR levels in restraint paradigm**. One-way ANOVA: F (3, 21) = 4.588 **(D)**; F (3, 22) = 0.9630, P = 0.4278 **(E)**; F (3, 22) = 11.79; P < 0.0001 **(F). (G-I) Semiquantitative analysis and representative immunoblot of p-ERK**_**1/2**_**/β-actin, ERK**_**1/2**_**/β-actin and p-ERK**_**1/2**_**/ERK**_**1/2**_ **levels in restraint paradigm**. One-way ANOVA: F (3, 19) = 2.94; P = 0.0494 **(G)**; F (3, 21) = 0.5587; P = 0.6481 **(H)**, F (3, 22) = 2,290; P = 0.1064 **(I)**. ^*^p < 0.05, ^**^p < 0.01 vs Cnt; ^+^p < 0.05, ^++^p < 0.01 vs Cnt R; ^#^p < 0.05, ^##^p < 0.05 vs SB24 R (Tukey’s test). **(J-L) Semiquantitative analysis and representative immunoblot of p-Akt/β-actin, Akt/β-actin and p-Akt/Akt levels in tail pinch paradigm**. One-way ANOVA: F (3, 23) = 0.7468; P = 0.5353 **(J)**; F (3, 23) = 0.3078; P = 0.8195 **(K)**; F (3, 24) = 0.9801; P = 0.4186 **(L). (M-O) Semiquantitative analysis and representative immunoblot of p-mTOR/β-actin, mTOR/β-actin and p-mTOR/mTOR levels in tail pinch paradigm in tail pinch paradigm**. One-way ANOVA: F (3, 24) = 0.4483; P = 0.7208 **(M)**; F (3, 24) = 0.6504; P = 0.5904 **(N)**; F (3, 24) = 1.067; P = 0.3817 **(O). (P-R) Semiquantitative analysis and representative immunoblot of p-ERK**_**1/2**_**/β-actin, ERK**_**1/2**_**/β-actin and p-ERK**_**1/2**_**/ERK**_**1/2**_ **levels in tail pinch paradigm**. One-way ANOVA: F (3, 22) = 0.8227; P = 0.4953 **(P)**; F (3, 23) = 2.744; P = 0.0663 **(Q)**; F (3, 23) = 6.016; P = 0.0035 **(R)**. ^+^p < 0.05 vs Cnt TP (Tukey’s test). Data are shown as mean ± S.E.M (n = 4 – 8 per group).

**Figure 4.**
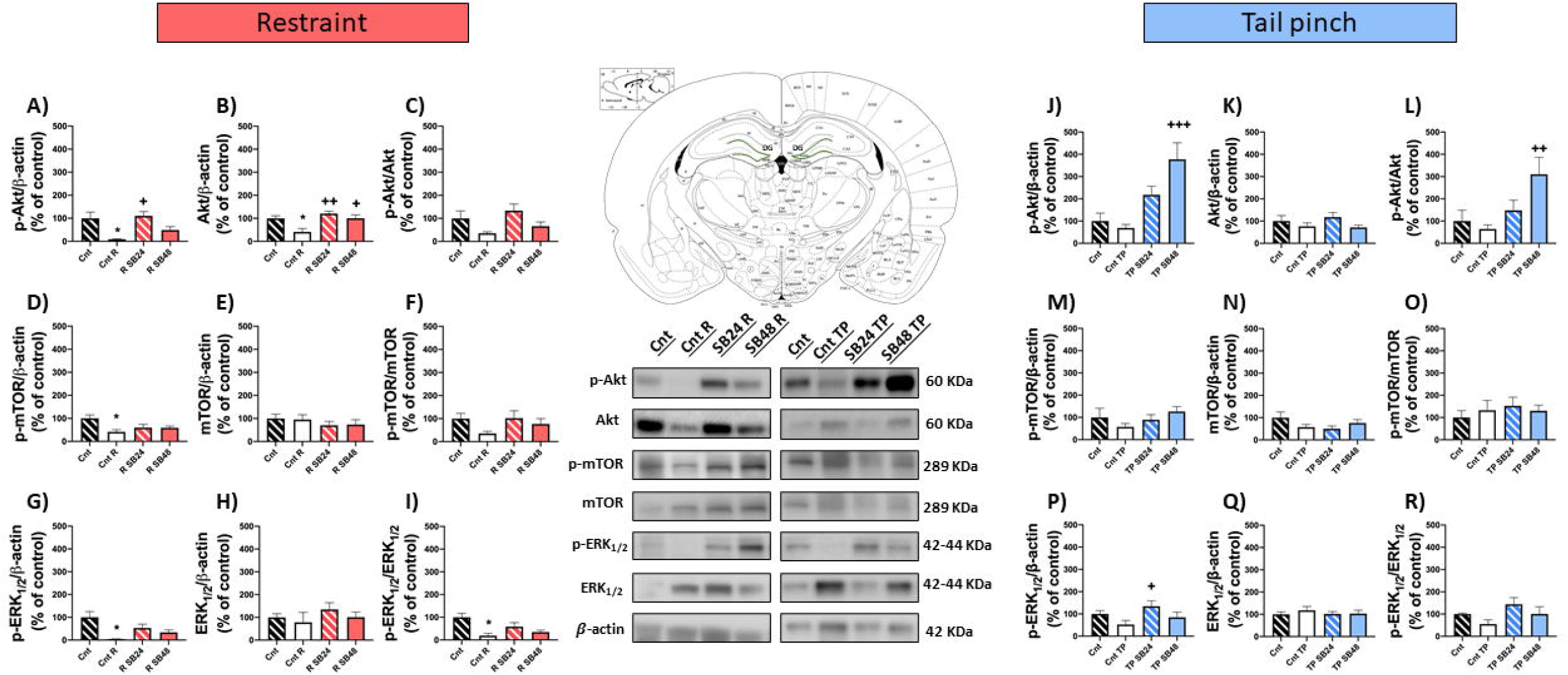
**Dentate gyrus analysis in restraint and tail pinch stress paradigms (A-C) Semiquantitative analysis and representative immunoblot of p-Akt/β-actin, Akt/β-actin and p-Akt/Akt levels in restraint paradigm**. One-way ANOVA: F (3, 24) = 3,838; P = 0.0224 **(A)**; F (3, 24) = 4.951; P= 0.0081 **(B)**; F (3, 24) = 2.086; P = 0.1287 **(C). (D-F) Semiquantitative analysis and representative immunoblot of p-mTOR/β-actin, mTOR/β-actin and p-mTOR/mTOR levels in restraint paradigm**. One-way ANOVA: F (3, 24) = 3.663; P = 0.0264 **(D)**; F (3, 24) = 0.5961; P = 0.6237 **(E)**; F (3, 23)= 0.9933; P = 0.4135 **(F). (G-I) Semiquantitative analysis and representative immunoblot of p-ERK**_**1/2**_**/β-actin, ERK**_**1/2**_**/β-actin and p-ERK**_**1/2**_**/ERK**_**1/2**_ **levels in restraint paradigm**. One-way ANOVA: F (3, 23) = 4.390; P = 0.0139 **(G)**; F (3, 24) =0.7024; P = 0.5599 **(H)**; F (3, 23) = 5.051; P = 0.0078 **(I)**. ^*^p < 0.05 vs Cnt; ^+^p < 0.05, ^++^p < 0.01 vs Cnt R (Tukey’s test). **(J-L) Semiquantitative analysis and representative immunoblot of p-Akt/β-actin, Akt/β-actin and p-Akt/Akt levels in tail pinch paradigm**. One-way ANOVA: F (3, 23) = 9.915; P = 0.0002 **(J)**; F (3, 23) = 1.442; P = 0.2563 **(K)**; F (3, 24) = 4.678; P = 0.0104 **(L). (M-O) Semiquantitative analysis and representative immunoblot of p-mTOR/β-actin, mTOR/β-actin and p-mTOR/mTOR levels in tail pinch paradigm**. One-way ANOVA: F (3, 24) = 1.513; P = 0.2366 **(M)**; F (3, 24) = 1.806; P = 0.1730 **(N)**; F (3, 22) = 0.2821; P = 0.8377 **(O). (P-R) Semiquantitative analysis and representative immunoblot of p-ERK**_**1/2**_**/β-actin, ERK**_**1/2**_**/β-actin and p-ERK**_**1/2**_**/ERK**_**1/2**_ **levels in tail pinch paradigm**. One-way ANOVA: F (3, 24) = 2.817; P = 0.0606 **(P)**; F (3, 24) = 0.3584; P = 0.7835 **(Q)**; F (3, 24) = 2.188; P = 0.1157 **(R)**. ^+^p < 0.05, ^++^p < 0.01, ^+++^p < 0.001 vs Cnt TP (Tukey’s test). Data are shown as mean ± S.E.M (n = 4 – 8 animals per group).

### 3.5 The dose of D3R antagonist is determinant for the activity of the Akt/mTOR and ERK_1/2_ pathways in the BLA and DG during the reactivation of cocaine-associated rewarding memories

The blockade of D3R had different effect on the activity of Akt, mTOR and ERK_1/2_ during the reinstatement of cocaine-CPP depending on the triggering stressor and the dose of antagonist administered. The BLA of animals that received 24 mg/kg of SB-277011-A prior being restrained to induce the relapse in the cocaine-seeking behaviour exhibited significantly higher p-Akt levels in comparison with mice that were injected with vehicle instead the D3R antagonist (Fig. 3A). In contrast, p-mTOR and p-ERK_1/2_ levels, despite higher than those of animals that received vehicle before the restraint episode, did not reach statistical significance (Fig. 3D,G). As total Akt, mTOR and ERK_1/2_ levels were not affected by the blockade of D3R (Fig. 3D,E,F), p-Akt/Akt, p-mTOR/mTOR and p-ERK_1/2_/ERK_1/2_ ratios in the BLA of mice that received the lower dose of the D3R antagonist prior restraint were enhanced, although only mTOR phosphorylation reached statistical significance with regard to animals that were injected with vehicle before the acute session of psychological stress. In contrast, the dose of 24 mg/kg of SB-211077-A during the reinstatement of cocaine-CPP induced by tail pinch did not modify p-Akt, p-mTOR and p-ERK_1/2_ levels (Fig. 3J,M,K) nor total Akt, mTOR and ERK_1/2_ levels (Fig. 3K,N,Q) in the BLA when compared with Cnt-TP group and, consequently, we did not detect significant alterations of the ratios p-Akt/Akt, p-mTOR/mTOR and p-ERK_1/2_/ERK_1/2_ in this area regarding animals that received vehicle instead the D3R blocker before the tail pinch session (Fig. 3L,O,R).

The DG of animals that were administered with 24 mg/kg of SB-277011-A before being restrained to induce the reinstatement of the cocaine-CPP exhibited significantly higher levels of p-Akt, but not p-mTOR nor p-ERK_1/2_, in comparison with mice that were injected with vehicle instead the D3R antagonist (Fig. 4A,D,G). Whereas no alterations were found in mTOR and ERK_1/2_ total levels, Akt levels were significantly higher in the DG of animals injected with 24 mg/kg of the antagonist in comparison with mice receiving vehicle prior the restraint episode (Fig. 4B,E,H). Despite that, p-Akt/Akt, p-mTOR/mTOR and p-ERK_1/2_/ERK_1/2_ in the DG of mice that received the lower dose of the D3R antagonist before being restrained exhibited a tendency to increase with regard to those receiving vehicle-injected before the stressful episode, although these enhancements were not statistically significant (Fig. 4C,F,I). On the other hand, after the administration of the lower dose of the D3R antagonist before the tail pinch episode were detected no significant differences in the levels of p-Akt and p-mTOR in comparison with mice injected with vehicle (Fig. 4M,P) in contrast to p-ERK_1/2_ levels, that were significantly augmented (Fig. 4JP). Total proteins levels in this region were not affected by the administration of 24 mg/kg of SB-277011-B (Fig. 4K,N,Q), and the resultant phosphorylation ratio of these kinases in the DG of mice that were injected with 24 mg/kg of SB-277011-B and subjected to a tail pinch session before the reinst test did not differ significantly of those of animals receiving vehicle instead the D3R antagonist (Fig. 4L,O,R).

In the BLA of mice that were injected with the higher dose of SB-277011-B before the restraint session p-Akt and p-mTOR levels were similar to those of vehicle-injected restrained animals and showed a trend to decrease regarding animals injected with 24 mg/kg of the D3R blocker prior being restrained (Fig. 3A,D) in contrast with p-ERK_1/2_ levels, that were significantly higher than those of animals that received vehicle before the restraint episode and similar to those of mice administered with the lower dose of the D3R antagonist prior the restraint session (Fig. 3G). Total Akt, mTOR and ERK_1/2_ remained unchanged (Fig. 3B,E,H), which resulted in a significant decrease of p-Akt/Akt and p-mTOR/mTOR in the BLA in comparison with animals injected with 24 mg/kg of SB-277011-B (Fig 3C,F), while p-ERK_1/2_/ERK_1/2_ were statistically similar in comparison with this group (Fig. 3I). In the DG of animals injected with the higher dose of SB-277011-B before being restrained we did not detect significant differences in the levels of p-Akt, p-mTOR and p-ERK_1/2_ with regard to animals that received vehicle or the lower dose of SB-277011-B prior the stressful stimulus, although p-Akt clearly tended to decrease when compared with mice that received the lower dose of the D3R antagonist (Fig. 4A,D,G). In contrast with total levels of Akt in the DG of mice injected with the higher dose of the D3R blocker prior restraint, that were significantly enhanced in comparison with those of animals injected with vehicle (Fig. 4B), total mTOR and ERK_1/2_ remained unaltered (Fig. 4E,H). As a result, we did not detect significant differences in the phosphorylation ratio of Akt, mTOR and ERK_1/2_ in the DG after the administration of 48 mg/kg of the D3R antagonist in comparison with those of mice that received the lower dose of SB-277011-B (Fig. 4C,F,I).

The administration of 48 mg/kg of SB-277011-B prior tail pinch did not modify p-Akt and p-mTOR (Fig. 3J,M) nor total Akt and mTOR protein levels (Fig. 3K,N) in the BLA. Therefore, the activity of both kinases remained unaltered (Fig. 3L,O) in this nucleus during the blockade of cocaine-CPP reactivation induced by a physiological stressor. As p-ERK_1/2_ levels showed a trend to augment and total ERK_1/2_ levels did not change in the BLA of mice subjected to tail pinch that were administered with the higher dose of the D3R antagonist with regard to those of animals receiving vehicle (Fig. 3P,Q), p-ERK_1/2_/ERK_1/2_ ratio was significantly enhanced with regard to vehicle-injected animals before the tail pinch session (Fig. 3R). In disparity, in the DG of mice injected with 48 mg/kg of SB-277011-B before an acute episode of physiological stress we did not find significant changes in p-mTOR and p-ERK_1/2_, (Fig. 4M,P) nor total ERK_1/2_ and mTOR levels (Fig. 4N,Q) in comparison with tail-pinched animals receiving vehicle or 24 mg/kg ofthe D3R antagonist and, thus, the activities of ERK_1/2_ and mTOR were unchanged (Fig. 4O,R). In opposition, p-Akt significantly augmented in the DG of animals injected with the higher dose of the D3R blocker and subjected to tail pinch with regard to tail pinched animals that received vehicle (Fig. 4J). As total Akt did not change (Fig. 4J), p-Akt/Akt statistically increased in the DG after receiving 48 mg/kg of the D3R antagonist to prevent the reactivation of cocaine-CPP induced by tail pinch (Fig. 4L).

### 3.6 Corticosterone plasma concentration might influence the activity of Akt/mTOR pathway but not ERK_1/2_ in the BLA and DG

We then studied, by means of Pearson’s correlation analysis, the possible relationship between the activity (as the ratio of phosphorylated/total protein levels) of Akt, mTOR and ERK_1/2_ in the BLA and the DG and the reinstatement of cocaine-seeking behaviours induced by psychological or physiological stress, but we did not find any significant correlations between those activities and the time spent by mice in the cocaine-paired compartment during the reins test nor the reinstatement score (data not showed). Next, we investigated whether the phosphorylation ratio of these kinases were related with the concentration of plasmatic corticosterone of animals subjected to acute psychological or physiological stress to induce the reactivation of cocaine-associated rewarding memories, and we detected high and positive correlations with p-Akt/Akt and p-mTOR/mTOR in the BLA of mice that were injected with 48 mg/kg of the D3R blocker and therefore did not relapse in cocaine-CPP after being restrained (Fig. 5A), with p-mTOR/mTOR in the BLA of mice that received 24 mg/kg of the D3R antagonist prior the psychological stressor (Fig. 5B) and with p-Akt/Akt in the DG of animals that were administered with 48 mg/kg of SB-277011-A before the same stimulus (Fig. 5C). Finally, we studied whether Akt, mTOR and ERK_1/2_ activities in the BLA and the DG correlated with the locomotor activity (measured as the difference in the number of total entries to both saline- and cocaine-paired compartments during the reinst test minus that during the ext test) of animals. We found that this difference of entries, on the one hand, significantly correlated highly and negatively with and p-mTOR/mTOR in the BLA of mice that received 48 mg/kg of the D3R antagonist before being restrained and, thus, did not reinstate the cocaine-CPP (Fig. 5D) and, on the other hand, significantly correlated positively with p-ERK_1/2_/ERK_1/2_ in the BLA of mice that were injected with the superior dose of SB-277011-B before the physiological stress episode and, consequently, did not reactivate cocaine-CPP (Fig. 5E).

**Figure 5.**
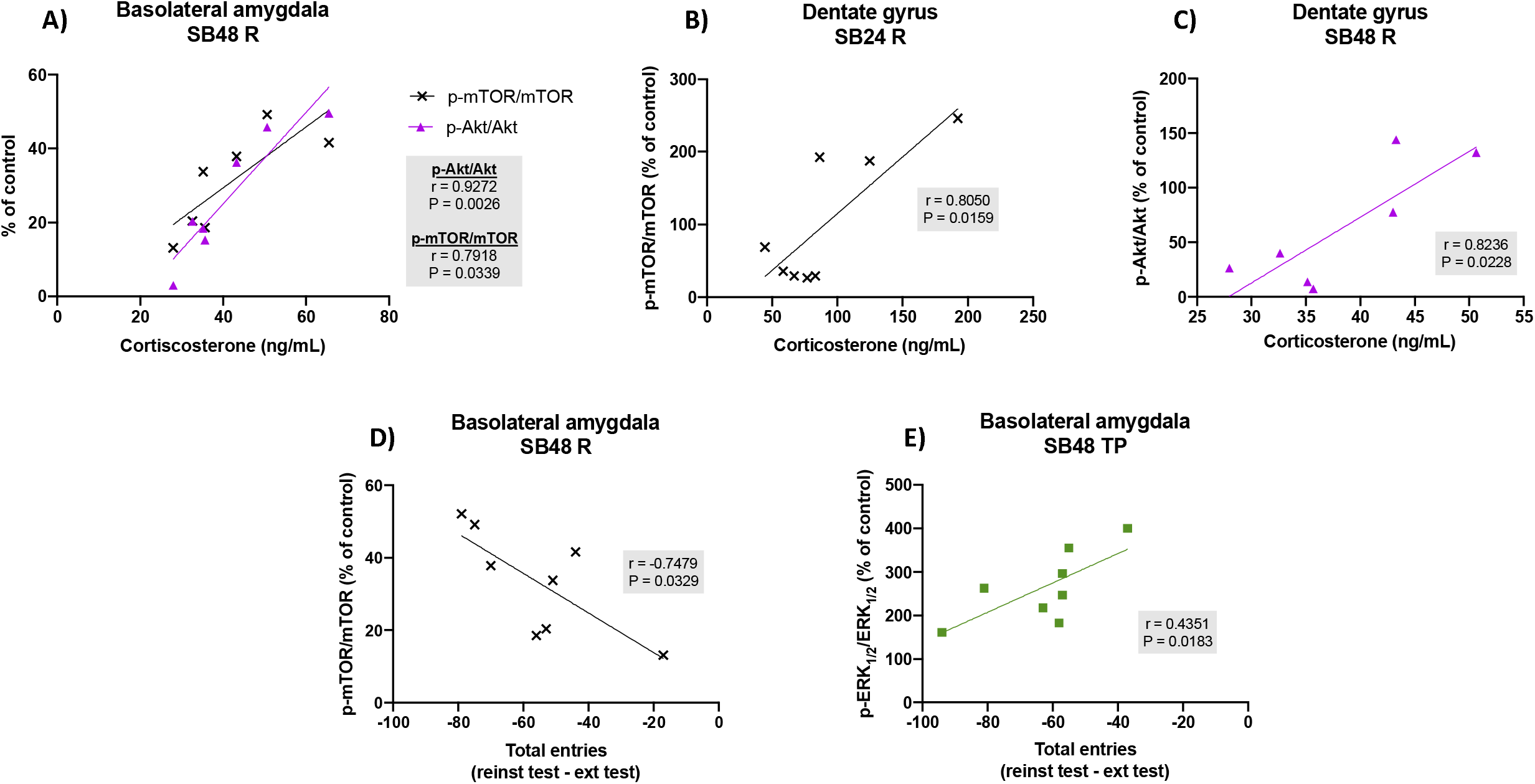
**(A) Correlation analysis of corticosterone in plasma of animals that experienced restraint and received the D3R antagonist at 48 mg/kg (SB48 R), p-Akt/Akt and p-mTOR/mTOR levels in basolateral amygdala. (B) Correlation analysis of corticosterone in plasma of animals that experienced restraint and received the D3R antagonist at 24 mg/kg (SB24 R) and p-mTOR/mTOR levels in dentate gyrus. (C) Correlation analysis of corticosterone in plasma of SB48 R mice and p-Akt/Akt levels in dentate gyrus. (D) Correlation analysis of total entries, expressed as the total entries during reinstatement (reinst) test minus those during extinction (ext) test, and p-mTOR/mTOR in basolateral amygdala of SB48 R animals. (E) Correlation analysis of total entries and p-ERK**_**1/2**_**/ERK**_**1/2**_ **levels in basolateral amygdala of animals that experienced tail pinch and received the SB-277011-A at 48 mg/kg (SB48 TP)**. Correlation analyses were performed through Pearson’s test (n = 4 – 8 animals per group).

## 4. Discussion

Concurrently with our previous investigations [6], this study shows that an episode of restraint or tail pinch reactivated the previously extinguished cocaine-associated rewarding memories responsible for the CPP. In addition, this work uncovers that the locomotor activity of animals restrained to induce the reinstatement of cocaine-CPP was diminished with regard to the extinction stage in contrast to mice tail-pinched, that displayed similar locomotor activity during the extinction and reinstatement tests. Besides the significant differences in their blood corticosterone levels, the disparity in the locomotor activity of animals depending upon the type of stress that triggered the reinstatement of cocaine-seeking behaviour might be as well related with the significant decreases in p-Akt, p-mTOR and p-ERK_1/2_ in the BLA and DG observed in animals restrained, that were not found when the relapse in cocaine-CPP was provoked by tail pinch. The present study also confirms our previous data [6] showing that the blockade of D3R prevented the reactivation of cocaine-seeking behaviours in mice subjected to an episode of psychological or physiological stress as well as the parallel enhanced corticosterone plasma concentration, and additionally reveals that the effect of D3R antagonism on the activity of Akt, mTOR and ERK_1/2_ in the BLA and DG is determined by the kind of stressor used to reinstate the cocaine-CPP. While the blockade of D3R prior tail pinch increased the activity of these kinases in the amygdalar and hippocampal areas analysed, the dose of antagonist administered seemed to influence the activity of the Akt/mTOR and MEK/ERK_1/2_ pathways when restraint triggered the relapse in cocaine-seeking behaviours. Corticosterone may be, in part, responsible for these variations as we found high correlations between its plasma levels and the activity of mTOR and/or Akt in both the BLA and DG of animals that were injected with the D3R antagonist before being restrained. Besides, our correlation analyses uncover that the locomotor activity of animals injected with the higher dose of the D3R antagonist, SB-277011-A, might be partly related with mTOR activity in the BLA when the reactivation of cocaine rewarding memories was induced by restraint, and on p-ERK_1/2_ phosphorylation ratio in the same brain region when the reinstatement of cocaine-CPP was provoked by an acute session of tail pinch.

Despite stress has been recognised as a risk factor for the relapse in drug-seeking behaviours for more than three decades, not all the stressors are able to induce the reinstatement of these conducts in different animal models for addiction research [5]. Whereas restraint was long known to induce the reactivation of cocaine-CPP [5], the present study confirms tail pinch as an effective stressor to reinstate this behaviour, what to our knowledge was first reported in our previous investigation [6]. Notwithstanding corticosterone is not essential for stress-induced reinstatement of drug seeking behaviours, it has been reported to have a permissive role [5]. Both psychological and physiological stressors are able to activate the HPA axis and eventually increase glucocorticoids plasma levels [9], which concurs with our data. Nevertheless, the higher corticosterone plasma concentration of restrained animals regarding those subjected to tail pinch might be due to the different anatomical substrates implicated in the psychological and physiological stress response [9]. Moreover, in discrepancy with mice restrained, corticosterone blood concentration of tail-pinched mice correlated highly with their reinstatement score. Altogether, these data seem to indicate a different degree of activation of the hypothalamic stress system depending upon the kind of stressor and might suggest that, once over certain plasma levels of glucocorticosteroids, this parameter does not influence cocaine-seeking behaviours. Nevertheless, corticosterone blood concentration does not seem to affect distinctly to the locomotor activity displayed during the relapse in cocaine-CPP by mice that were subjected to tail pinch and those restrained. Our study also shows that mice receiving vehicle before being restrained did augment the time spent in the cocaine paired-chamber in parallel with a decrease in the number of entries to the same compartment when compared with the extinction test, in discrepancy to mice that reactivated CPP by physiological stress, that increased the time spent in the drug-associated compartment but did not variate the number of entries to the cocaine-paired chamber regarding the extinction test. This might suggest that the drug-associated context gains a more powerful emotional value after the reinstatement of CPP by restraint than by tail pinch, which might be related as well with the distinct neurosystems involved in the response to stress depending on the type of stimulus [9].

The contexts associated with addictive substances can aberrantly activate the learning and memory brain systems, thus prompting drug-directed behaviours even after prolonged abstinence periods [30]. In drug-induced CPP paradigm, the rewarding effects of the addictive compound are associated by pavlovian conditioning with a specific environment, and the reactivation of this preference provokes the recall of the pleasurable memories paired with the drug [30, 31]. The retrieval of distinct kinds of memories has been associated with alterations in the activity of Akt/mTOR and MEK/ERK_1/2_ pathways in several brain areas [22, 25], although the manner in which they change remains controversial. Classically, the activity of ERK_1/2_ and Akt/mTOR pathways, for their role in synaptoplasticity, had been considered vital for hippocampal declarative memory retrieval [32]. Nonetheless, ours and others’ studies recently reported diminished mTOR and/or Akt phosphorylation after the recall of drug-associated aversive and rewarding memories in the NAcc, hippocampus and BLA [7, 8, 33, 34]. The present investigation uncovers a different pattern of activation of these kinases in the BLA and DG determined by the kind of stressor that triggered the recall of cocaine-associated memories. While the physiological stressors act, at least partially, by means of defined receptor systems, the psychological or psychosocial stressful stimuli are processed through less specific exteroceptors and/or somatic inputs with emotional components [9]. Additionally, the neurocircuitries integrating the response to each kind of stress has been reported to be different [35]. The effects of tail pinch have been shown to be mediated by dopaminergic and orexinergic neurotransmission systems and to influence cognitive processes [36–39]. Nevertheless, these effects have been reported to be integrated in the ventral hippocampus and medial amygdala [38, 40], which might explain the lack of activity observed in the DG and BLA during the reinstatement of cocaine-CPP induced by this stressor. On the other hand, the changes observed during the reactivation of the cocaine-seeking behaviour induced by restraint were concordant with those observed in these brain regions when the recall of rewarding memories of the drug was provoked by a psychosocial stressor [8].

Dopaminergic inputs from the VTA to its projection areas are vital for the reinstatement of drug use [41]. In particular, D3R has a critical role in the maintenance of the behavioural outcomes of drug addiction [16]. Accordingly, and in agreement with the data reported in this study, it was published before by ours and others’ laboratories that the blockade of D3R prevented the relapse in abused drugs-seeking behaviours provoked by stress [6–8, 17, 18, 42–44]. The BLA receives direct dopaminergic innervation from the VTA, as does the hippocampal DG [11–13]. In addition, we have detected before D3R expression in both areas and shown that its blockade diminished the availability of these receptors during the reinstatement of cocaine-CPP induced by psychosocial stress, as occurred also in the NAcc [7, 8]. Consequently, the activity of D3R-coupled biochemical pathways might be modified after the antagonism of this receptor in order to impede the relapse in cocaine-CPP, as it has been published before by us and others [7, 8, 21]. Present results reflect different effects of D3R antagonism on Akt, mTOR and ERK_1/2_ during the reinstatement of cocaine-CPP depending upon the kind of stressor. When the reactivation of cocaine rewarding memories was induced by tail pinch, the administration of SB-277011-A induced an increase of the activity of Akt and ERK_1/2_ that seems dose-dependent for ERK_1/2_ in the BLA and for Akt in the DG, with no apparent alteration of mTOR phosphorylation in none of the areas analysed. Akt phosphorylation at Ser 473 results in its full activation and, as a consequence, to lead to the phosphorylation of its downstream substrates, being mTORC1 among them [45]. The unchanged levels of p-mTOR in both the BLA and DG despite Akt phosphorylation at Ser 473 in animals that received the D3R blocker before the tail pinch session might respond to the complex modulation of the Akt/mTOR signalling pathway, that has been published to be composed by more than 250 components and 478 links between them to participate in a number of cellular events [46]. The MEK/ERK_1/2_ signal has been suggested to cooperate with Akt in some cellular events and compensate the last’s function when it is deactivated [47]. Hence, our results might indicate that D3R, through the balanced modulation of ERK_1/2_ and Akt activity, participates in the reactivation of cocaine-rewarding memories provoked by physiological stress.

In divergence, our data indicate a more intricated participation of D3R in the reinstatement of cocaine-seeking behaviours evoked by psychological stress as the lower dose of SB-277011-A, that was ineffective to prevent the reinstatement of cocaine-CPP, produced a general increase of the phosphorylation and activity of Akt, mTOR and ERK_1/2_ in both the BLA and DG, whereas the administration of 48 mg/kg of the D3R antagonist, that efficiently blocked the restraint-evoked reactivation of the cocaine-seeking behaviour, affected distinctly to the activity of these kinases in both the amygdalar and hippocampal areas analysed. It seems illogical that 48 mg/kg of SB-277011-B had no effect on the activity of mTOR or Akt in the BLA after the reinstatement of cocaine CPP when the lower dose tested in this investigation altered their phosphorylation ratio. Nonetheless, other psychodrugs such as tricyclic antidepressants are known to exert their effects in a restricted therapeutical window and to have no efficiency when administered at lower or higher doses [48]. We propose that the ability of D3R to prevent the reactivation of cocaine-rewarding memories responsible for the relapse in CPP triggered by psychological stress results from the balanced modulation of Akt/mTOR and MEK/ERK_1/2_ pathways achieved with the higher dose of its antagonist, and that this equilibrium would differ in both the hippocampal and amygdalar areas implicated in memory retrieval analysed in this work. Whilst in the BLA the decrease in the activation of Akt/mTOR pathway appears to be compensated by higher ERK_1/2_ phosphorylation ratio, present data does not allow us to suggest a clear regulation of Akt/mTOR and MEK/ERK_1/2_ pathways in the DG.

Since no correlations were found between the phosphorylation ratio of these kinases and the reinstatement score of restraint- or tail pinch-induced cocaine-CPP, it seems that their activities cannot be considered as indicators of relapse in drug-seeking behaviour. However, the blood levels of corticosterone after the blockade of D3R to avoid the relapse in cocaine-CPP evoked by psychological stress might be relevantly related with the activity of mTOR and Akt in the BLA and DG as their positive correlations ranged from very high to almost perfect, although the mechanism responsible for this link remains yet to be deciphered. In addition, and as reported before [6], the augmented corticosterone blood concentration induced by the reactivation of cocaine-CPP triggered by both psychological and physiological stress was prevented by both 24 and 48 mg/kg of SB-277011-B in what seems a dose-dependent manner. Nevertheless, only the higher dose of the D3R blocker was able to antagonise the reinstatement of this behaviour. These data might indicate that the dopaminergic neurons in the mesolimbic system would be strongly activated and in consequence would release high levels of DA in their projection areas, therefore being needed a high dose of the D3R blocker to prevent the stress-induced reactivation of cocaine-CPP and supporting, then, a facilitating but not decisive function of glucocorticoids in the relapse in drug-seeking.

The results reported in this work reveals that the administration of SB-277011-A provoked a decrease of the locomotor activity during the reinstatement of tail pinch-induced cocaine-CPP regarding extinction test, which might be suggested to be due to D2 receptors rather than the D3R-selective antagonism. Nonetheless, SB-277011-A (up to 90 mg/kg, PO) and other selective D3R blockers failed to affect spontaneous or stimulant-induced locomotion and did not provoke catalepsy at a dose of two times the higher administered in the present study, in opposition to D2 antagonists [49]. Moreover, SB-277011-A has 80-to 100-fold selectivity over other DA receptor and high affinity for human (pKi 7.95) and rat (pKi 7.97) D3R [50]. We also found a moderate positive correlation of the locomotor activity of tail-pinched mice with the phosphorylation ratio of ERK_1/2_ in the BLA when the higher dose of the antagonist was administered to prevent the reinstatement of cocaine-CPP the moderately correlated. In addition, and regardless of no differences in the locomotor activity during the restraint-induced reactivation of CPP after the administration of SB-277011-A, this locomotion highly correlated negatively with mTOR activity in the BLA when 48 mg/kg of the antagonist was administered. Yet, what appears an attenuating effect of SB-277011-A on the locomotor activity during the psychological and physiological stress-induced reactivation of cocaine rewarding memories and the underlying mechanisms cannot be easily interpreted on the basis of the present data.

In conclusion, this study strengths the hypothesis of a critical function of D3R, by means of a complex and balanced regulation of Akt/mTOR and MEK/ERK_1/2_ pathways in brain nuclei implicated in emotional and contextual memory consolidation and recall, in the retrieval of drug memories, thus supporting a promising therapeutic potential of its antagonists to prevent the stress-induced reinstatement of drug-seeking behaviours. Our work will, therefore, aid to minimise the relapses in drug use. Nevertheless, our investigation leaves unresolved questions such as the mechanism by which D3R regulates glucocorticoid release or locomotor activity during relapse in cocaine use, that remain to be elucidated in future investigations.

## Credit authorship contribution statement

A.F.-G.: Investigation, Visualization, Data Curation and Writing—Original Draft. R.G.-B.: Investigation, Visualization, Data Curation. V.G.-M.: Investigation, Visualization, Data Curation and Writing—Original Draft. J.M.H.: Investigation and Resources. M.V.M.: Conceptualization, Validation, Formal analysis, Data Curation, Funding acquisition, Project administration, Supervision, and Writing—review & editing. C.N.: Conceptualization, Validation, Formal analysis, Data Curation, Funding acquisition, Project administration, Supervision, and Writing—Original Draft. All authors have read and agreed to the published version of the manuscript.

## Acknowledgements

This study was financially supported by MCIN/AEI/10.13039/501100011033 and by “ERDF A way of making Europe” (grants SAF2017-85679-R and PID2020-113557RB-I00), and by Fundación Séneca, Región de Murcia, Spain (grant 21133/SF/19). Aurelio Franco-García is granted by “Ayuda para la Formación de Profesorado Universitario” program of MICINN (FPU19/01722).

## Data availability statement

The data in current study can be obtained from the corresponding authors upon reasonable demand.

## Author Contributions

M.Victoria Milanés and Cristina Núñez were responsible for the conception and design of the experiment as well as writing and reviewing the final version of the manuscript. Rocío Guerrrero-Bautista performed all the behavioural studies. Juana M. Hidalgo processed all the brain samples. Aurelio Franco-García and Victoria Gómez-Murcia contributed to the experiments, the writing of the manuscript and preparing the figures.

